# Tumor Segmentation using temporal Independent Component Analysis for DCE-MRI

**DOI:** 10.1101/2022.12.16.520830

**Authors:** Joonsang Lee, Qun Zhao, Marc Kent, Simon Platt

## Abstract

In this study, we developed the temporal independent component analysis (tICA) to solve the partial volume effect (PVE) in canine brain tumor segmentation. The performance of the tICA is compared to that of spatial ICA (sICA) and an expert manual delineation of tumors based on three criteria: percent volume overlap or Dice coefficient, and percent volume difference. Seven in vivo DCE-MRI datasets of spontaneously occurring canine brain tumors were acquired on a 3 Tesla MRI system. The mean value of percent volume overlap (76.00%) between sICA and manual segmentation is lower than that between tICA and manual segmentation, which is 81.11%. In conclusion, the performance of the two ICA methods for segmenting tumors is very close to that of the expert delineation method. However, the tICA has the advantage over the sICA method in its ability to separate independent tissue signals in a voxel containing different types of tissues.

## Introduction

Dynamic Contrast-enhanced magnetic resonance imaging (DCE-MRI) is widely used in a cancer-imaging tool. It is a noninvasive, clinical imaging technique that involves the study of tumor angiogenesis^1^ and in the development and trial assessment of antiangiogenic and vascular disrupting compounds^2^. Also, the studies of DCE-MRI includes, but are not limited to, noninvasive assessment of tumor microenvironment^3^, predictors of clinical outcomes including treatment response to chemotherapy^4,5^, detection of rheumatoid arthritis^6,7^, differentiation of tumor histopathology^8,9^ and analysis of the pharmacokinetic parameters^10^. With a high sensitivity of MRI, it has been widely applied to improve tumor detection and diagnosis^11^. In DCE-MRI, malignant tumors show faster and higher enhancement than normal tissue. In general, no enhancement curves or slow sustained enhancement curves are typically associated with normal or benign tissues. Rapid initial and stable late enhancement curves can be classified as suspicious, and rapid initial and decreasing late enhancement curves can be classified as malignant.

The tissue classification and anatomical segmentation are increasingly studied through MRI. Due to a growing the amount of MRI data, the automated method is required to develop accurate and reliable image analysis for classifying image regions. As a result, many automated computer-aided methods are proposed such as the region-growing method to segment lesions^12^, automated segmentation methods based on artificial intelligence techniques^13^, segmentation based on statistical pattern recognition techniques^14^, a semiautomatic algorithm based on the fuzzy c-means clustering^15^, a user-interaction-threshold method to extract the region of interest (ROI)^16^, automatic segmentation combining label propagation with decision fusion^17^, and a detection of deviations from normal brains using a multi-layer Markov random field framework^18^. Recently, independent component analysis (ICA) has been introduced to the field of DCE-MRI for the detection and characterization of breast lesions^19^, and identification of breast lesions as separate hemodynamic sources^20^.

In DCE-MRI, it requires repeated acquisition of T_1_-weighted images of a particular tissue of interest (TOI) before, during and after an intravenous administration of a bolus of a paramagnetic contrast agent (CA), typically a low-molecular weight gadolinium (Gd) compound. The contrast uptake curves in a TOI or voxels are often fitted using a pharmacokinetic model to give physiological information about blood flow, capillary leakage and related physiological parameters. In general, tumor tissues show a high and fast contrast uptake due to abundance of angiogenic microvessels in tumor tissues and normal tissues or benign tissues show no enhancement curves or slow sustained enhancement curves. In the studies of the detection and classification of tumor on DCE-MRI data, the independent component analysis (ICA) methods were recently used to identify breast tumor^20^. Also, application of ICA on DCE-MRI includes calculation of intravascular signal^21^, removing undersampling artifacts^22^, and assessment of cerebral blood perfusion^23^.

Independent component analysis (ICA), a special case of blind source separation (BSS) algorithms that separates a set of signals from a set of mixed signals without information of the source signals, finds the independent components by using statistical independence^24^. Applications of ICA have been found in many areas such as electrical recordings of brain activity as given by an electroencephalogram (EEG) device, analysis of functional MRI (fMRI), feature extraction, and medical image processing^25–27^. During the last decade, ICA becomes one of the most widely used statistical and computational method that can separate a multivariate signal into the original source signals by assuming that components are both statistically independent and nongaussian. Several different algorithms have been proposed from different approach such as the maximum likelihood estimation (MLE) that is a method of estimating the parameters of a statistical model, nonlinear PCA (NLPCA) algorithm that is the nonlinear equivalent of classical PCA, and reduces the observed variables to a number of uncorrelated principal component developed by Karhunen and Joutsensale (1994) and Oja (1995), the information maximization algorithm (infomax algorithm) by Bell and Sejnowski^28^, Joint approximate diagonalization of eigen-matrices (JADE) that is equivalent to informatics approaches and employs approximate joint diagonalization of fourth-order cumulant matrices proposed by Cardoso and Souloumiac^29^, and FastICA that is an efficient and popular algorithm based on fixed-point iteration and maximizing non-Gaussianity proposed by Aapo Hyvärinen^30^. The ICA methods can be divided into two groups, sICA and tICA. The sICA method finds a set of mutually independent spatial images such as tumor or normal tissue^31^, while tICA finds a set of independent time courses such as enhancing time curves in DCE-MRI^28,32–34.^

In the studies of tumor segmentation using MR data, partial volume effect (PVE) is one of the major difficulties and may result in inaccurate segmentation results due to inherent low spatial resolution of images^35^. The PVE occurs when more than one tissue type presents in a voxel and it blurs the intensity distinction at the border of two tissues such as the tumor and normal tissues. In an earlier study, sICA has been used to solve the PVE on large blood vessel^36^ and arterial input function^37^ in microPET.

In this study, seven in vivo DCE-MRI datasets from a canine model of spontaneously occurring brain tumors were acquired and analyzed with ICA. There are two complementary ways in ICA to decompose signals into original source signals, sICA and tICA. To resolve this difficulty presented by PVE in the segmentation of the canine brain tumor, this study uses tICA, which intends to separate two intrinsic tissues at the border. Although ICA has been used to discriminate source signals from biological mixture signals for DCE-MRI data^21^, to the best of our knowledge tICA has never been implemented in the segmentation of DCE-MRI. The tICA method was also compared with the sICA and the manual delineation of the lesions by an expert was taken as a reference standard in evaluating the methods.

## Material and Methods

ICA is a statistical and computational technique that extracts individual source signals from the measured mixture signals by means of statistical independence of the non-Gaussian source signals. A simple form of the ICA problem can be expressed by the following linear model:

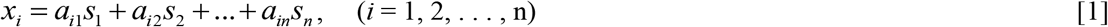

where *s_1_*, …, *s_n_* are the original source signals, *a_ij_* (*j* = 1,…,*n*) are weighting coefficients, and *x_i_* is the mixture signal which is weighted sums of the *sj*. In this basic ICA model, we assume that each mixture *x_i_* and each independent component *s_i_* is a random variable. To simplify the method and algorithm without loss of generality, we can perform centering the observable variables by subtracting the sample mean so that both the mixture variables and the independent components have zero mean. Then a whitening procedure was performed as preprocessing step using principal component analysis (PCA) and eigenvalue decomposition (EVD). The purpose of whitening is to make the components uncorrelated and their variances to be unity^33^. For the convenience, we can rewrite this model into vector-matrix notation

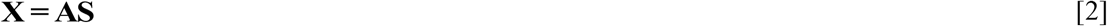

where **X** is the *m* × *n* measured data matrix and both **A** and **S** are unknown, and need to be estimated using ICA under the assumption that the component *s_j_* are statistically independent. We can obtain the independent component simply by

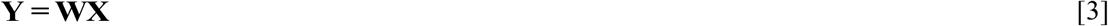

where **Y** is the estimated independent source signals and W is the unmixing matrix, and Y can be estimated by setting up a cost function which either maximizes the nongaussianity of the calculated *s_j_* or minimizes the mutual information. In this study, we have used the fastICA method, which use a fixed-point algorithm and the negentropy as a cost function to estimate the original signals.

### MRI data acquisition

Seven spontaneous occurring brain tumors were imaged using a transmit/receive knee coil on a 3.0 Tesla GE SIGNA HDX MR scanner (GE Medical Systems, Milwaukee, WI). The paramagnetic contrast agent, gadopentetate dimeglumine or Gd-DTPA, was injected intravenously as a bolus (0.2 mMol/kg) after the first acquisition pulse. The DCE-MRI protocol employed a standard T_1_-weighted, 2-D gradient refocused echo sequence to obtain dynamic serial images with the following parameters: TR of 34 ms, TE of 2.78 ms, 35° flip angle, acquisition matrix size of 192×192, field of view (FOV) of 182.25 cm^2^, a total of 5 slices, slice thickness of 3 mm, and NEX=1, scan time of 5.9 minutes (a total of 41 acquisitions and a temporal resolution of 8.7 seconds).

### Manual segmentations by expert

Among the 41 acquired DCEMRI images, the most clearly enhanced tumor image on a single slice was selected for the manual segmentation. An expert (MK, an experienced neurologist) manually traced the outline of the seven brain lesions. The size of image is 256 × 256 with a 256-gray level. In this study, the manual delineation was used as a standard reference and compared with that of tICA and the sICA, respectively.

### Segmentations using spatial ICA

In this study, we have selected the whole brain region as a mixture and assumed that there are three spatially independent components: normal brain tissue, tumor tissue, and noise. A three-source example of Eq. (1) can be rewritten as

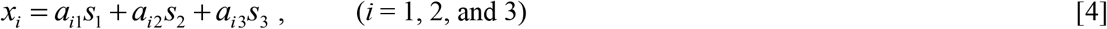

where *s_1_, s_2_*, and *S_3_* are spatially independent source components, respectively. The measured mixture data *x_1_, x_2_*, and *x_3_* were sampled from the dynamic series at three different points in time. First, the preprocessing was performed to make *x_i_* a zero-mean variable by centering *x_i_*. Whitening was performed as a preprocessing by transforming the observed vector *x_i_* linearly so that its components are uncorrelated and their variances to be unity. Then, each frame of the mixture images, *x_i_* (*i* = 1, 2, and 3), was converted into a 1D row vector.

The estimated independent component can be obtained by iteratively updating the unmixing matrix **W**. Each row vector *y_i_*, is then reformed into 2D image to construct the independent component map. To differentiate between normal brain tissue and tumor tissue, we calculated standard deviations, σ, of the background signals randomly selected from *x_1_* (i.e., independent component map of tumor), and then we removed this background signals by selecting the 3σ as the threshold of the background signals based on the three-sigma rule (^38^), which indicates that most of the signals (about 99.7%) of the background lie within three standard deviations of the mean for a normal distribution.

### Segmentations using a temporal ICA

Regarding MR data, brain tissue segmentation is usually complicated by PVE because a voxel may contain two tissue types. DCE-MRI data is measured over a period of time, where a dynamic contrast-enhanced time curve is acquired from each voxel of the image. It is assumed that the observed time curve signal is a linear mixture of different source signals, e.g., tumor tissue and normal brain tissue. In a partial volume voxel, the signal intensity can be determined by the following linear mixture

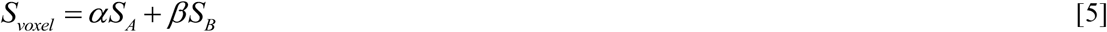

where α and β are weighted coefficient, and *S_A_* and *S_B_* represent signal intensity of tissue A and tissue B, respectively.

In the tICA of the DCE-MRI data, we have selected a region covering brain tumor as a ROI, and assumed that there are two independent components: tumor tissue and background signal that is normal brain tissue. A two-source example of Eq. [1] can be rewritten as

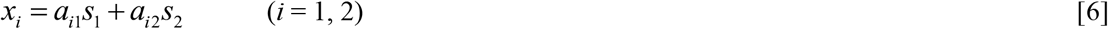

where *s_1_* and *s_2_* are temporally independent source components, respectively. The measured mixture data *x_1_* was sampled from a voxel in TOI and *x_2_* was sampled from each voxel in entire image.

The estimated independent component **Y** can be obtained by iteratively updating the inverse of the mixing matrix **W = A^-1^**, also known as the unmixing matrix by means of statistical independent properties. We have set *y_1_* as a TOI signal and *y_2_* as a normal brain tissue signal. A mixing matrix **A** was then converted from the unmixing matrix **W** and then obtained the *a_11_* map, where *a_11_* is the weighting coefficient of *s_1_* from each voxel in the frame.

The segmentation threshold was determined based on the binary mask map created from comparing the weighting coefficient ratio between *a_11_* and *a_12_*. If the weighting coefficient ratio *a_11_*/*a_12_* of a certain voxel is greater than one, indicating the tumor tissue takes more than 50% volume of the voxel, then that voxel was assigned the value of one as TOI; otherwise the ratio less than 1 implies normal tissue is more than 50% volume of the voxel, then a zero is assigned representing a normal brain tissue. The final segmented TOI mask was calculated by maximizing the Pearson’s correlation coefficient between the tumor sizes from an *a_11_* map and a binary mask map. In this way, an incorrect assignment of the voxel at the background when the weighting coefficient ratio is close to one will be removed based on the threshold that is obtained from the Pearson’s correlation coefficient with *a_11_* map.

### Comparison

The segmented area estimated by the tICA was compared with the area extracted by the sICA. Comparison was performed based on three methods: percent volume overlap or Dice’s coefficient as defined in Eq. [7], and percent volume difference as defined in Eq. [8]^39,40^.

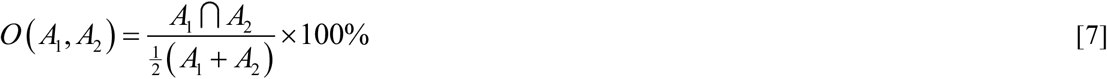

where ***A***_**1**_ □ ***A***_**2**_ means an overlapped area between two regions from each method. The maximum value of 100 represents that they overlap perfectly. It is noted that the overlap coefficient between two different areas can be slightly decreased by the spatial location shifts. We have also calculated area difference between two regions, which is insensitive to spatial shift.

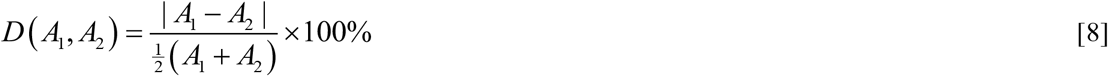

For the identical volume size, *D*(*A_1_,A_2_*) will have the value of zero indicating that there is no difference between two area in volume.

## Results

Segmentation of the lesion volumes using sICA and tICA were compared to an expert’s manual delineation. Figure 1 represents the results of the manual, sICA, and tICA segmentation, respectively, for all seven canine objects used in this study. The first column represents the manual segmentation and the second and third columns represent the segmentations from sICA and tICA, respectively. It is noted that tICA generated two original sources, tumor (s_1_) and normal tissue (s_2_), as indicated in Eq. [6]; meanwhile the sICA presented three original sources, tumor (s_1_), normal tissue (s_2_), and noise (s_3_), as shown by Eq. [4].

**Figure 1.**
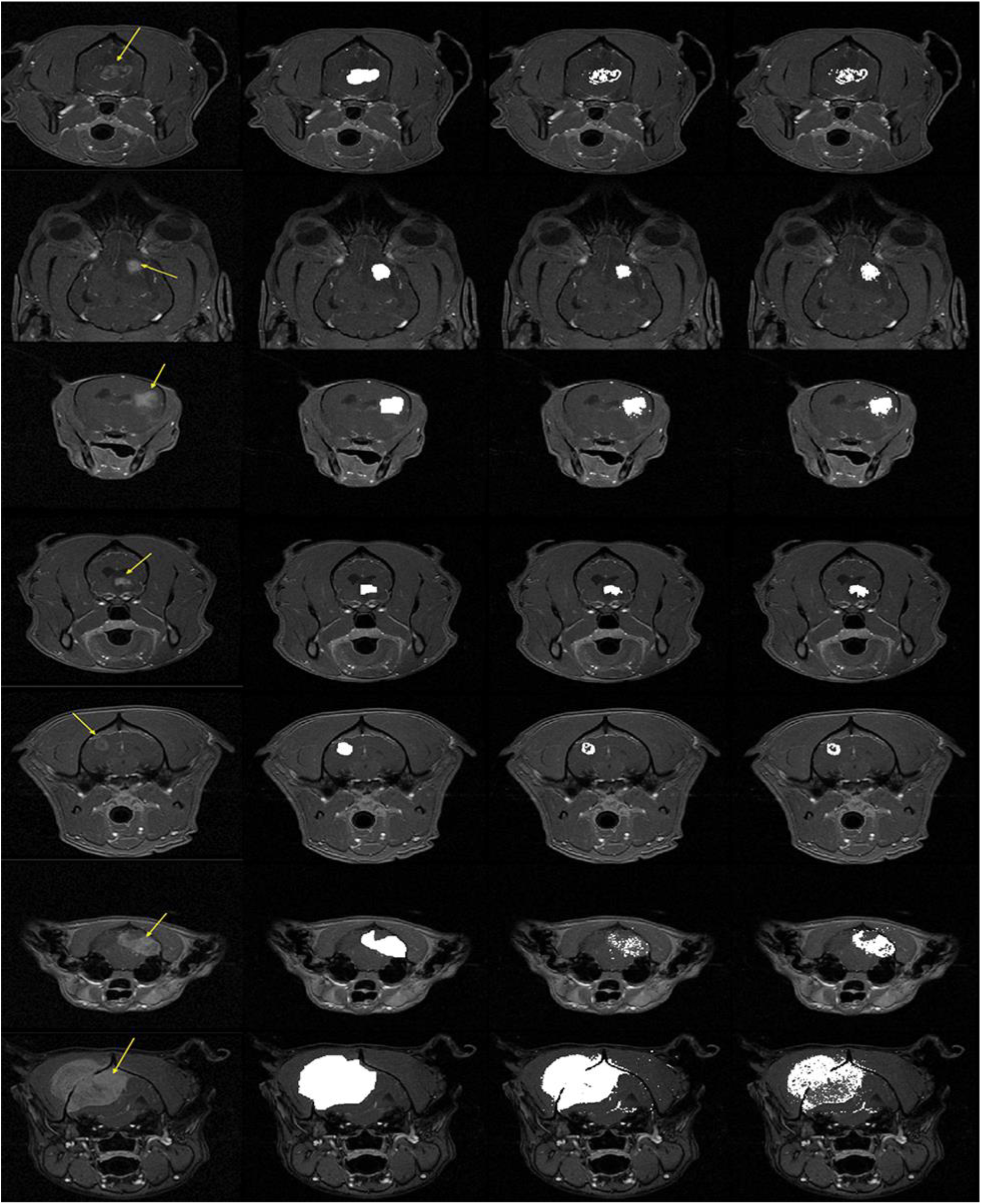
The results of each segmentation method:(1^st^ column) all 7 canine brain tumor images (tumors were indicated with arrows), (2^nd^ column) expert’s delineation segmentation, (3^rd^ column) sICA segmentation, and (last column) tICA segmentation.

Figure 2 presents a closer look at the dataset 1by showing the original canine brain tumor image (a), detailed view of a small frame (covered by a box in (a)) that contains tumor and surrounding brain tissues (b), the *a_11_* map that was obtained from the weighting coefficient of the tumor signal for each voxel obtained from tICA (c), and a binary map of the weighting coefficient ratio, *a_11_/a_12_* that is bigger than 1 (d). In the binary map (d), incorrect assignments of voxels at the background were also seen (isolated spots). After removing these incorrectly assigned voxels, a final segmented binary mask is shown in Figure 2(e). However, due to inhomogeneity nature of the tumor, this final binary mask is not a smooth and continuous map, which has a low percent volume overlap (72.10%) and a high percent volume difference (45.39%) with the manual segmentation. To further investigate the inhomogeneity appearance of the tumor, figure 2(f) represents an *a_11_* profile across the center of the TOI, along the line shown in 2(c). The *a_11_* profile showed that high *a_11_* values correlate well with enhancing tumor area, while low *a_11_* values correlate with non-enhancing brain tissues.

**Figure 2.**
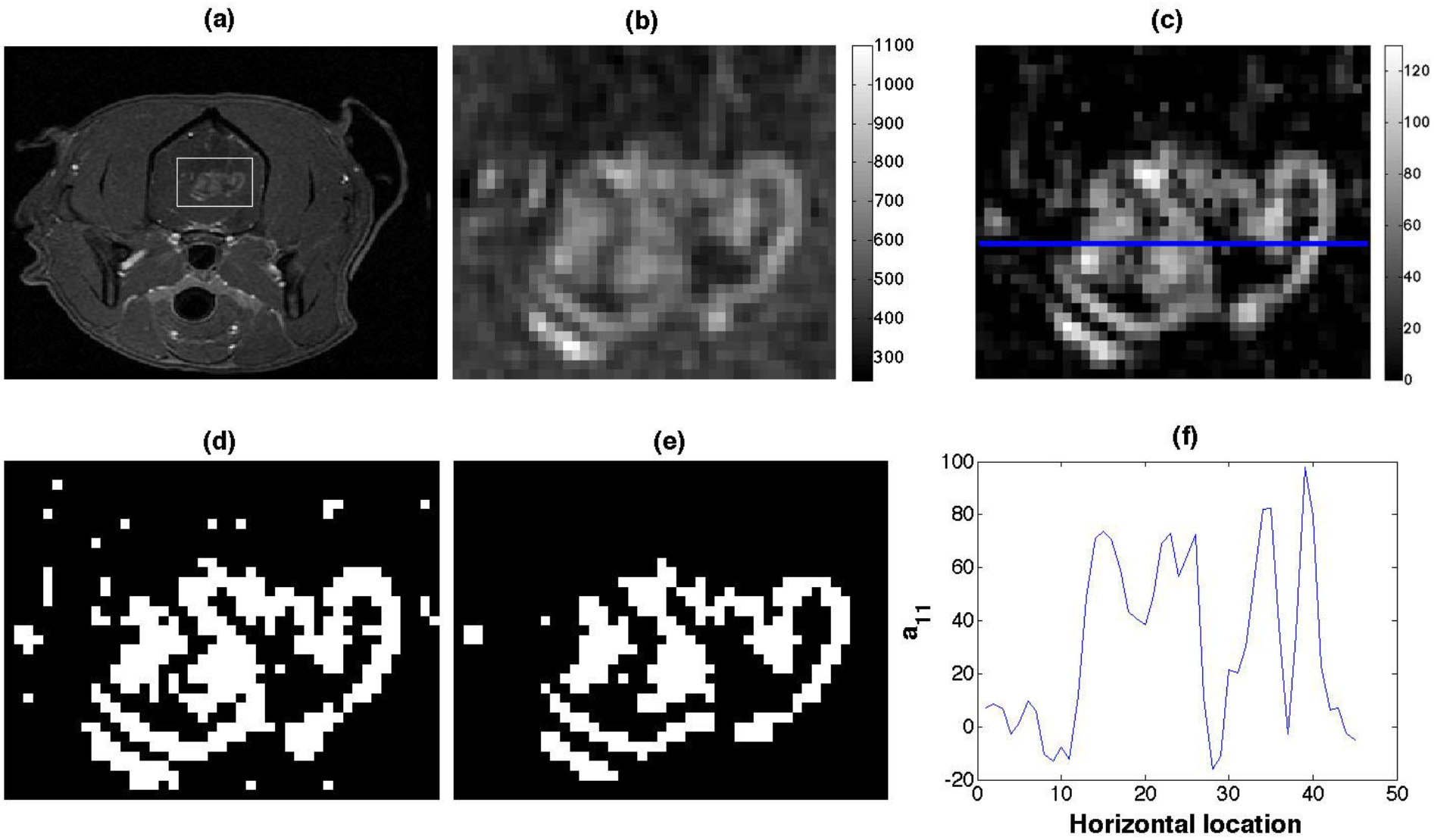
A closer look at the dataset 1: (a) Original canine brain tumor image. (b) A small frame that contains tumor and normal brain tissue. (c) *a_11_* map from temporal ICA. (d) Binary map from the weighting coefficient ratio, *a_11_/a_12_*, which is bigger than 1. (e) Final segmented mask from temporal ICA. (f) *a_11_* profile plot at the horizontal center of TOI showing that the values of the coefficient of tumor tissue *a_11_* have high values and low values at the normal brain tissues.

Figure 3 presents closer view of two homogeneous tumors, datasets 2 and 3, as shown in (a) and (d). The *a_11_* profiles across the horizontal and vertical enhancing tumor area of dataset 2 and dataset 3 are shown in Figure 3(b) and 3(c), 3(e) and 3(f), respectively. Both tumor cases show relatively high percent volume overlaps (90.45% and 89.68%) and relatively low percent volume differences (2.39% and 6.47%), respectively, with the manual segmentation. It is seen that the values of *a_11_* across the border between tumor and surrounding brain tissues increase or decrease smoothly indicating that the PVE possibly occurs along the border of tumors. By comparing coefficients of each source (tumor or normal brain tissue), the tumor borderline can be easily determined from the coefficient ratio *a_11_/a_12_* of each voxel. Measured sizes of each segmented area are shown in Table 1.

**Figure 3.**
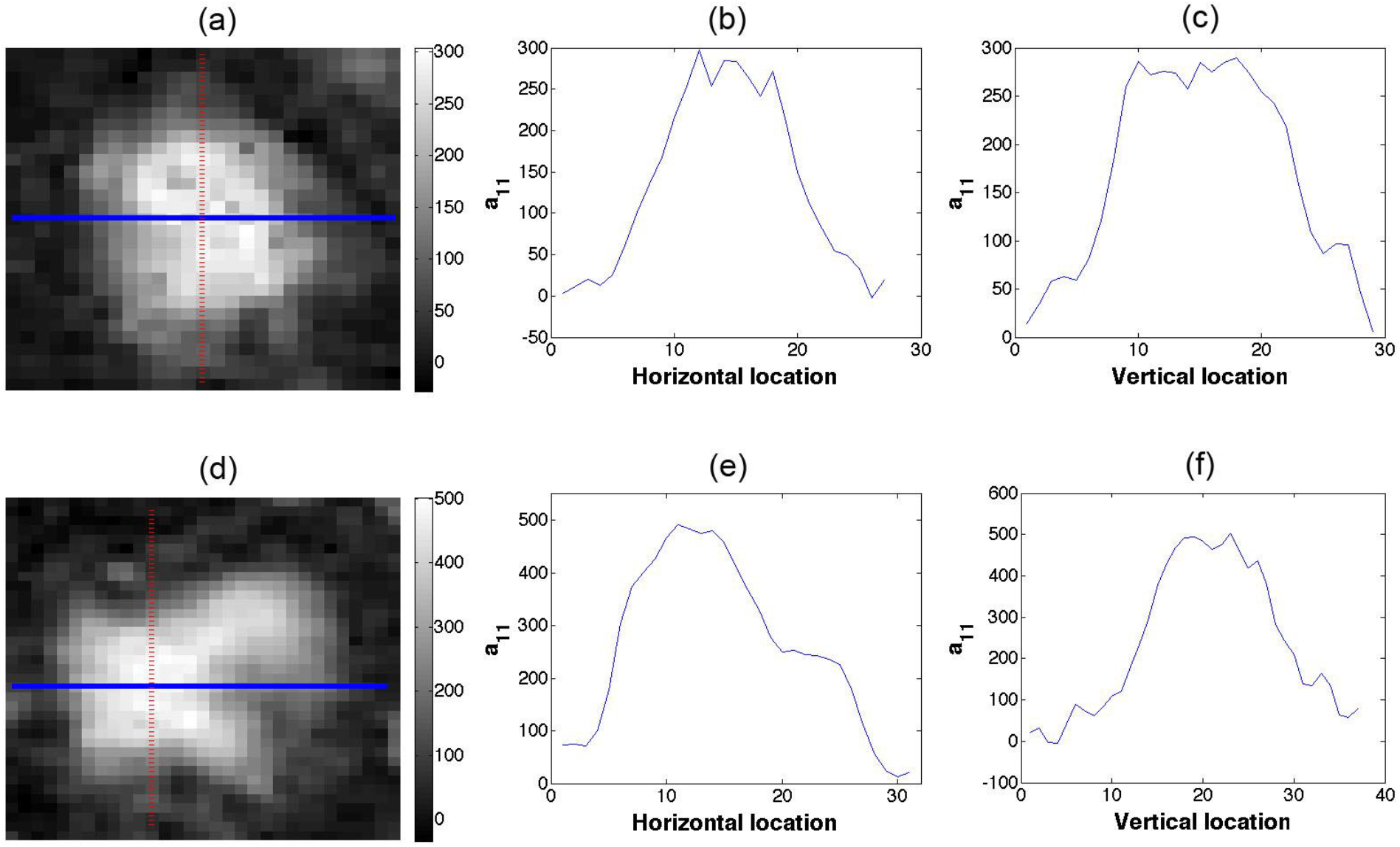
A closer look at the dataset 2 and dataset 3: *a_11_* profile plots at the horizontal and vertical cross section of TOI for dataset 2 (a~c) and dataset 3 (d~f). It is shown that the values of *a_11_* at the border between tumor and normal brain tissues increase or decrease gradually indicating that the PVE occurs at the border of tumors.

**Table 1.** Segmented Area.

| Datasets | Segmentation Method |  |  |
| --- | --- | --- | --- |
| | Spatial ICA ($cm^2$) | Temporal ICA ($cm^2$) | Manual ($cm^2$) |
| Data 1 | 1.74 | 1.76 | 2.79 |
| Data 2 | 0.66 | 1.13 | 1.01 |
| Data 3 | 1.04 | 1.02 | 1.09 |
| Data 4 | 0.92 | 1.02 | 0.96 |
| Data 5 | 0.68 | 0.71 | 1.18 |
| Data 6 | 0.63 | 2.70 | 2.23 |
| Data 7 | 17.84 | 11.62 | 17.47 |

The results of each method were also compared using two indexes, the percent volume overlap or Dice’s coefficient (Eq. [7]) and the percent volume difference between two regions (Eq. [8]). Table 2 summarizes these results, including the mean value and the standard deviation. The mean values of percent volume overlap and percent volume difference between sICA and manual segmentation were 76.00% and 38.07%, respectively. The mean values of percent volume overlap and percent volume difference between temporal ICA and manual segmentation were 81.11% and 24.84%, respectively. Between the two ICA segmentation methods, the percent volume overlap and percent volume difference were 79.44% and 29.94%, respectively. According to these results, the overlap volume percentage between tICA and manual segmentation has a slightly higher percentage rate than that between sICA and manual segmentation. Meanwhile, the percent volume difference between tICA and manual segmentation has a smaller percentage rate than that between sICA and manual segmentation.

**Table 2.** Overlap (%) and Difference (%) between three methods sICA and tICA represent spatial and temporal ICA, respectively.

|  | sICA and Manual |  | tICA and Manual |  | sICA and tICA |  |
| --- | --- | --- | --- | --- | --- | --- |
|  | O(A<sub>1</sub>,A<sub>2</sub>) | D(A<sub>1</sub>,A<sub>2</sub>) | O(A<sub>1</sub>,A<sub>2</sub>) | D(A<sub>1</sub>,A<sub>2</sub>) | O(A<sub>1</sub>,A<sub>2</sub>) | D(A<sub>1</sub>,A<sub>2</sub>) |
| Data 1 | 74.15 | 46.46 | 72.10 | 45.39 | 88.42 | 1.13 |
| Data 2 | 77.64 | 42.58 | 90.45 | 2.39 | 76.59 | 40.29 |
| Data 3 | 90.27 | 4.97 | 89.68 | 6.47 | 93.05 | 1.50 |
| Data 4 | 85.04 | 3.67 | 81.25 | 13.46 | 90.46 | 17.11 |
| Data 5 | 72.73 | 54.55 | 77.44 | 44.10 | 86.11 | 11.11 |
| Data 6 | 40.55 | 112.14 | 83.62 | 21.83 | 44.82 | 96.19 |
| Data 7 | 91.58 | 2.10 | 73.25 | 40.23 | 76.60 | 42.24 |
| Mean & STD | 76.00±17.33 | 38.07±39.63 | 81.11±7.34 | 24.84±18.30 | 79.44±16.57 | 29.94±33.68 |

## Discussion

In this study, sICA and tICA methods were used for the segmentation of canine brain tumors using spatial and temporal independency of tumor characteristics shown in the DCE-MRI data. For application of the ICA in DCE-MRI, sICA was commonly used to detect and classify tissue types by assuming the spatial independence of these tissues. In this paper, we have used the 3σ rule of the background signals as a threshold for segmentation in the sICA. For tICA, the segmentation threshold was determined by weighting coefficients of the mixing matrix. The difference of the two methods is that sICA can provide spatial information by separating the DCE-MRI data into physiologically meaningful components, while tICA can provide temporal information from the corresponding time courses. In addition, tICA is potentially capable of solving the problem of PVE, one of the major difficulties in tumor segmentation. The difficulty with PVE is that it blurs the intensity distinction at the border of the tumor and surrounding tissues. In a voxel with PVE (i.e., two tissues reside in one voxel), the MR signal results from both tissues and signal intensity of this voxel can be expressed as the sum of the signal from each tissue, as shown in the PVE model (Eq. [5]). In principle, ICA can separate this mixed signal into individual tissue signals, if they are statistically independent, to resolve this PVE. If the two signals do not meet the statistical independency condition, this method will likely fail.

One drawback of the tICA method for segmentation is that one has to identify the actual number of source signals in advance. In this work, we have selected a small frame that only contains two different tissue types, i.e., tumor and non-tumor tissues, so that we can limit the number of source signals to two for the feasibility of the tICA segmentation. If one needs to further differentiate non-tumor tissues, the number of source, *i*, should be increased in Eq. [6]. In this case, the size of the unmixing matrix will be *i × i* and this could make segmentation more difficult. The reason for this is that if the enhancing curves of tumor and muscle tissues look similar to each other then there might be some correlation between them, implying the statistical independency condition may not be met. Figure 4 shows such an example, where Figure 4(a) presents three randomly selected, representative enhancing curves from tumor, muscle, and normal brain tissues. Figure 4(b) shows the scatter plot of one representative independent components S_1_ (enhancing curve from tumor) and S_2_ (enhancing curve from normal brain tissue), which has a small correlation coefficient (0.08±0.07); while the scatter plot of the independent component S_1_ and S_3_ (enhancing curve from muscle tissue) shown in Figure 4(c) has a high correlation coefficient (0.85±0.03). Five voxels were selected from each tissue (tumor, muscle, and normal brain tissue), respectively, from data 2, which is the representative homogeneous tumor case. Mean values and standard deviations of correlation coefficients are calculated from 25 possible pairwise combinations and listed in Table 3.

**Figure 4.**
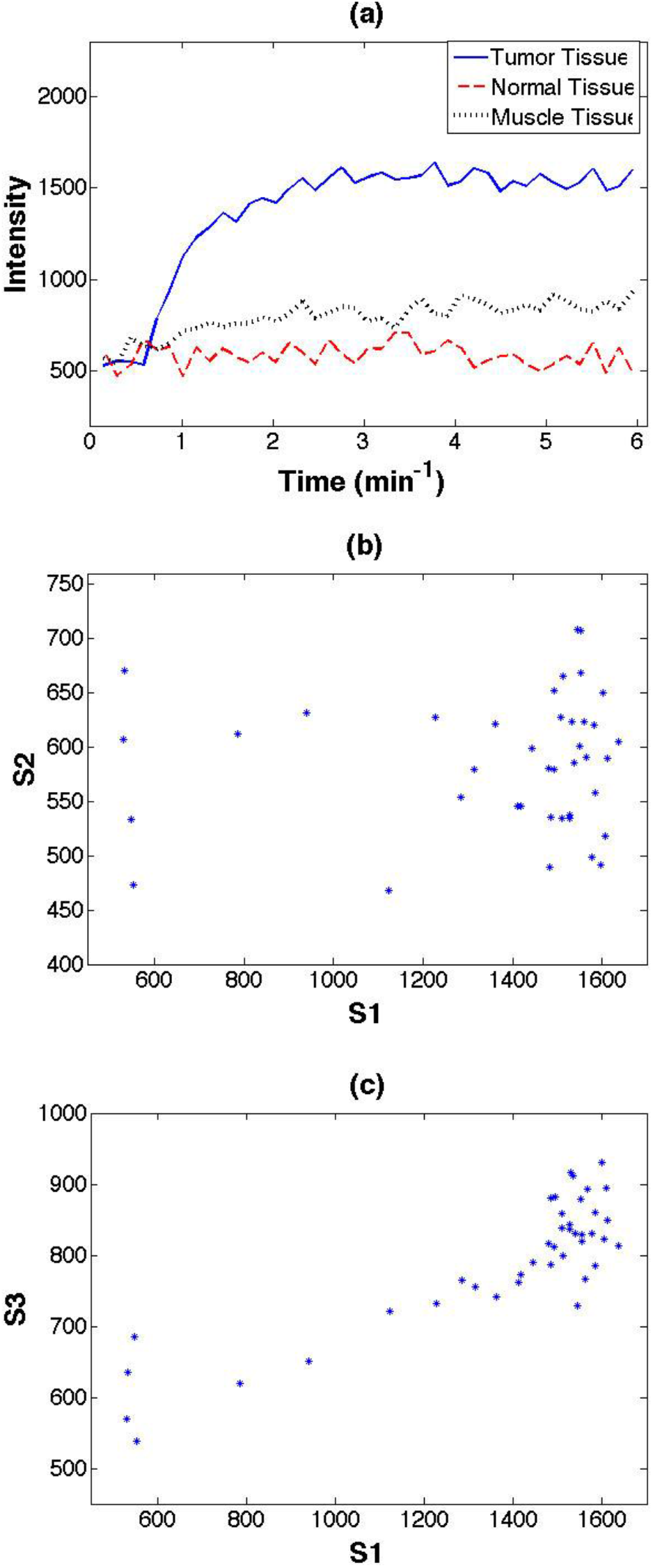
(a) Three enhancing curves from tumor, muscle, and normal brain tissues. (b) The scatter plot of the independent components S1 (tumor) and S2 (normal brain tissue). (c) The scatter plot of the independent components S1 and S3 (muscle tissue).

**Table 3.**
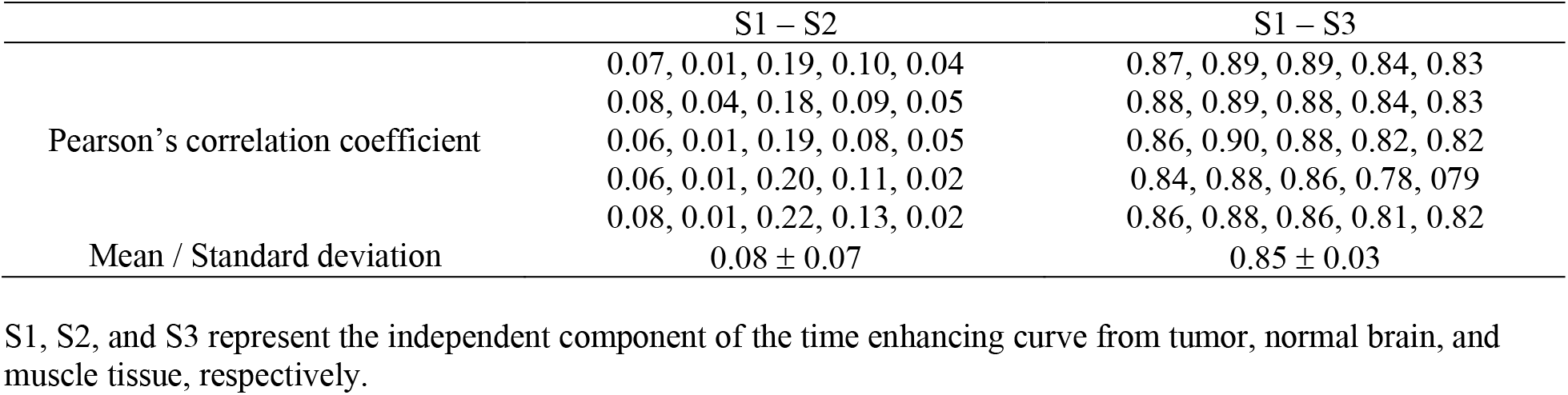
Pearson’s correlation coefficient between two enhancing time curves.

|  | S1 – S2 | S1 – S3 |
| --- | --- | --- |
| Pearson's correlation coefficient | 0.07, 0.01, 0.19, 0.10, 0.04 | 0.87, 0.89, 0.89, 0.84, 0.83 |
|  | 0.08, 0.04, 0.18, 0.09, 0.05 | 0.88, 0.89, 0.88, 0.84, 0.83 |
|  | 0.06, 0.01, 0.19, 0.08, 0.05 | 0.86, 0.90, 0.88, 0.82, 0.82 |
|  | 0.06, 0.01, 0.20, 0.11, 0.02 | 0.84, 0.88, 0.86, 0.78, 0.79 |
|  | 0.08, 0.01, 0.22, 0.13, 0.02 | 0.86, 0.88, 0.86, 0.81, 0.82 |
| Mean / Standard deviation | 0.08 ± 0.07 | 0.85 ± 0.03 |

S1, S2, and S3 represent the independent component of the time enhancing curve from tumor, normal brain, and muscle tissue, respectively.

In summary, we have applied the temporal and sICA method for the segmentation of the brain tumor using the DCE-MRI data. These two ICA methods were compared with an expert’s manual delineation, which was used as a standard reference. We have shown that the performance of two ICA methods for segmenting tumor is very close to that of the expert’s delineation method. For the two ICA methods, we have shown that the tICA has the advantage over the sICA method in its ability to separate independent tissue signals in a voxel containing different types of tissues automatically by solving a PVE while sICA needs to set a threshold manually.

## Acknowledgements

We would like to thank all of the staff in the BioImaging Research Center (BIRC) at The University of Georgia for their contribution to this study.

## Data Availability

The datasets generated during and/or analyzed during the current study are available from the corresponding author on reasonable request.

## Author Contributions

Project conception and design were by J.L. and Q.Z. The data collection and preprocessing were performed by J.L., M.K., and S.P. The software programming, statistical analysis, and interpretation were performed by J.L. The manuscript was written by J.L. and all authors reviewed the manuscript.

